# Heterosis in crosses between remnant populations of the rare prairie forb *Silene regia*: implications for restoration genetics

**DOI:** 10.1101/2024.10.13.618093

**Authors:** Isabelle A. Turner, Juan Diego Rojas-Gutierrez, Bob Easter, Christopher G. Oakley

## Abstract

**Premise:** Calls to adopt seed sourcing strategies in biological restoration that preserve local adaptation while maximizing genetic variation (e.g., regional admixture provenancing) are increasingly common. Heterosis, the increased fitness of progeny from between-relative to within-population crosses, could provide an added benefit under such strategies, but several open questions remain.

**Methods:** We quantified heterosis in crosses between three small remnant populations of the rare prairie forb *Silene regia*. We measured early fitness components (seed number per fruit, germination, and juvenile survival) in a greenhouse. Adult fitness components (survival and reproduction) were quantified over two flowering seasons in two different environments: a field experiment simulating the initial stages of a restoration, and a greenhouse. Our approach is unique in estimating heterosis in an environment most relevant to the early stages of a restoration and in comparing heterosis under field and controlled conditions.

**Key Results:** The consequences of between-population crosses for cumulative fitness in the field were strongly positive in two of the populations (281% and 50% heterosis) and nearly neutral in a third. Heterosis was generally stronger when measured under field conditions, and stage specific patterns of heterosis varied among populations.

**Conclusions:** Regional admixture provenancing in restorations should be beneficial for this species. We advocate for more research on heterosis in rare species, particularly in restorations. We also provide a cautionary note on how to calculate and report results for heterosis studies.

## INTRODUCTION

Limited genetic variation needed to adapt to variable or changing conditions, and the negative fitness consequences of fixed deleterious recessive alleles, are major issues in conservation and restoration genetics (Frankham, 2005; Willi et al., 2006; Bell et al., 2019). One way to probe the relative extent of deleterious recessive alleles fixed within populations in nature is heterosis, the increased fitness in between-population relative to within-population crosses (Fenster and Galloway, 2000; Paland and Schmid, 2003; Oakley and Winn, 2012; Lohr and Haag, 2015). Heterosis is common in crosses between natural populations (Fenster, 1991; van Treuren et al., 1993; Ouborg and Van Treuren, 1994; Luijten et al., 2002; Willi and Fischer, 2005; Pickup et al., 2012; Willi, 2013; Weisenberger et al., 2014; Koski et al., 2019; Oakley et al., 2019; Perrier et al., 2020; Söderquist et al., 2020; Thoen et al., 2025). Greater heterosis in small/less diverse compared to large/more diverse populations (Paland and Schmid, 2003; Escobar et al., 2008; Oakley and Winn, 2012; Lohr and Haag, 2015; Spigler et al., 2017) and in selfing compared to outcrossing populations (Busch, 2006; Oakley et al., 2015b) is consistent with a role of drift in fixing partially recessive deleterious alleles (Charlesworth, 2018). This suggests that heterosis may be particularly relevant in conservation and restoration of small remnant populations.

One context in which heterosis could be potentially useful is in restoration projects involving rare species. Restoration practitioners use seeds, or sometimes seedling plugs, to establish populations (Shaw et al., 2020), often including rare or threatened species that are dispersal- or seed-limited (Pywell et al., 2002; Seabloom et al., 2003; Clark et al., 2007). The method of seed sourcing and the genetics of the source populations can greatly influence establishment success (De Vitis et al., 2022). Traditionally it was thought that seeds should be sourced as locally as possible to minimize the risk of disrupting adaptation to local environmental (extrinsic) conditions or outbreeding depression due to (intrinsic) genetic incompatibilities (McKay et al., 2005; Gann et al., 2019; Török et al., 2024). However, potential seed sources for restoration are often small remnant populations from which only a small number of individuals (sometimes only a single individual) are sampled (Godefroid et al., 2011). Thus, the founding population may harbor fixed deleterious alleles and have limited genetic variation to adapt to fluctuating or changing conditions, particularly in the face of climate change. Because of these risks, many authors have called for some version of seed sourcing strategies that increase genetic variation while also minimizing outbreeding depression and the loss of local adaptation (Broadhurst et al., 2008; Godefroid et al., 2011; Breed et al., 2013; Bucharova et al., 2019; Rossetto et al., 2019; St. Clair et al., 2020; Hoffmann et al., 2021; Stojanova et al., 2021; Jordan et al., 2023; Fahey et al., 2025). One proposed strategy is “regional admixture provenancing” (Bucharova et al., 2019), where seed is sourced from multiple populations regionally to increase genetic variation. These populations would however come from similar habitat in the same eco-region, likely connected by gene flow prior to fragmentation, to minimize the risks of maladaptation and genetic incompatibilities. Additionally, heterosis following admixture in a restoration may provide at least a short term fitness boost that may aid establishment of rare species, analogous to the “catapult effect” proposed for the role of heterosis in the establishment of invasive species (Drake, 2006; Li et al., 2018). Despite the potential benefits, there are few empirical tests of the regional admixture provenancing approach (but see St. Clair et al., 2020) or the potential for heterosis during a newly established restoration.

Some outstanding questions in the study of heterosis more broadly are how the magnitude depends on experimental assay conditions and to what extent it varies across the life cycle. The former is somewhat surprising given the considerable work on similar questions in inbreeding depression which also involves changes in the level of expression of partially recessive deleterious alleles. For example, work by Dudash (1990) demonstrated greater inbreeding depression in field compared to greenhouse conditions, and since then the environmental dependence of inbreeding depression has been well documented (reviewed in Fox and Reed, 2010; Cheptou and Donohue, 2011). Heterosis is often estimated in the greenhouse, though studies estimating heterosis on plants transplanted into the field for at least part of the life cycle are becoming more common (Fenster, 1991; Ouborg and Van Treuren, 1994; Fenster and Galloway, 2000; Luijten et al., 2002; Wagenius et al., 2010; Maschinski et al., 2013; Weisenberger et al., 2014; Oakley et al., 2015b; Spigler et al., 2017; Oakley et al., 2019; Thoen et al., 2025). However, we are unaware of any comparison of heterosis in field vs. controlled conditions, though there are some comparisons in multiple environments (Fenster and Galloway, 2000; Willi et al., 2007; Li et al., 2018; Stojanova et al., 2021; Perrier et al., 2022). Unlike inbreeding depression which can be influenced by both strongly deleterious and more mildly deleterious alleles (Charlesworth and Willis, 2009), heterosis is expected to be mostly caused by the complementation of mildly deleterious alleles (Whitlock et al., 2000; Glémin et al., 2003; Charlesworth, 2018). Heterosis may therefore be weak for individual fitness components, particularly early in the life cycle (c.f., Husband and Schemske, 1996; Perrier et al., 2020). Therefore, obtaining accurate estimates of heterosis may require estimates of fitness encompassing as many life stages as possible.

Here we used within- and between- population crosses of three small remnant populations of a rare prairie forb, *Silene regia* Sims., to quantify heterosis for multiple fitness components in both a greenhouse experiment and a field experiment mimicking a newly established restoration. We addressed the following questions: 1) What is the magnitude of heterosis (or conversely, outbreeding depression) under restoration conditions in the field, and does it vary among populations? 2) Is the magnitude of heterosis greater in the field compared to controlled greenhouse conditions? and 3) How does the magnitude of heterosis vary across different life history stages? We expected to observe greater heterosis under field compared to greenhouse conditions, as well as greater heterosis in estimates of fitness incorporating later life history stages.

## MATERIALS AND METHODS

### Study system

*Silene regia* (royal catchfly) is a tap-rooted perennial in the Caryophyllaceae (Dolan, 1994). It is endemic to a dozen states in eastern North America and is thought to have been broadly distributed across prairies (Menges, 1991) prior to the widespread conversion of this habitat to agriculture in the 19^th^ century. Contemporary populations across much of the native range are small and isolated and exist as remnants in prairie cemeteries and along railroad rights-of-way and roadsides (Dolan, 1994). Remnant populations of *S. regia* are potentially at risk of genetic depletion (Menges and Dolan, 1998). This species does not vegetatively reproduce and achieves sexual reproduction via hummingbird pollination. It is self-compatible and some geitonogamous selfing may occur, but it is likely mostly outcrossing as the individual flowers are protandrous (Dolan, 1994; Menges, 1995).

We selected three remnant focal populations in different counties in western Indiana (Tip = Tippecanoe; Fou = Fountain; Ver = Vermillion). Exact locations of these populations are not provided because this species is critically imperiled in Indiana (NatureServe, 2024), but these are distinct populations, all separated by at least 14 km. These three populations include all known existing remnants within a 55 km radius, and there may be as few as 10 extant remnant populations in the entire state of Indiana (B. Easter, pers. obs.). While including more populations could have broadened our scope of inference, we restricted the sampling to remnants from the same local area as would be advised under regional admixture provenancing. The smallest population by census size was Tip (10 flowering individuals) from a ∼0.5 ha remnant, which has likely been small since the 19^th^ century (B. Easter, pers. obs.). The largest population was Ver (∼50 flowering individuals) from a ∼0.4 ha remnant. Population size reductions are more recent at this site where decades of regular mowing during the 20^th^ century up until the 1980’s restricted flowering individuals to fencerows (B. Easter, pers. obs.). The size of Fou, from a ∼0.4 ha remnant, was intermediate to the other populations (∼20 flowering individuals). While there is some variation in census size among these remnants, they are all quite small and are thus all expected to have limited genetic variation and harbor fixed deleterious mutations.

### Generation of seeds for the fitness assays

Bulk seed was collected from each population in September 2019. Our goal was to characterize population level heterosis, and the small number of flowering individuals in these populations precluded experimental designs accounting for distinct maternal lineages. We collected seeds from most of the reproductive individuals in the two smaller populations and from a haphazard subset of approximately 20 individuals in the largest population, Ver.

In early October 2019, seeds from each population were surface sterilized in a 10% bleach solution for 10 minutes, followed by three rinses in deionized-autoclaved water. The seeds were then placed into moist paper towels in bags and cold stratified in the dark at 4°C for 10, 30, or 60 days. Different stratification periods were used because 60 days would be ideal for germination, but starting the plants sooner was preferrable due to logistical constraints. Because of low overall germination we included individual maternal plants arising from all three seed stratification treatments. While there was a significant effect of seed stratification treatment (for seeds that produced the parental plants) on seed number per fruit following controlled crosses, this treatment (involving the generation prior to the experimental assay) did not qualitatively affect the results for either the effects of cross type or the interaction between maternal population and cross type (compare Table S1 to Table 1) so we omitted this term from subsequent analyses.

**Table 1.** ANOVA results for early fitness components of offspring derived from hand pollinations within and between three populations (Fou, Tip, and Ver) of *Silene regia* from western Indiana. Fixed effects include maternal and paternal populations, their interaction, and block (where appropriate, otherwise n/a). Models for both seed number per fruit and percent germination (per cell) employed normal error distributions (denominator degrees of freedom in parentheses). The model for juvenile survival employed a binomial error distribution. Contrasts between cross types are only reported for models with a *P* value < 0.1 for either the interaction between maternal population and paternal population, or both main effects of maternal and paternal populations, otherwise these contrasts are n/a. For the contrasts, WIN = within-population cross, BET = between-population cross, and the subscript pool indicates that the contrast involves all individuals of a given cross type (averaged over all three populations). Contrasts that start with a population code followed by a colon test for heterosis or outbreeding depression separately by maternal population.

|  | Seed number per fruit<br>(Denom. DF = 229) |  | Germination<br>(Denom. DF = 346) |  | Juvenile Survival |  |
| --- | --- | --- | --- | --- | --- | --- |
| | *F* | *P* | *F* | *P* | $\chi^2$ | *P* |
| Maternal population | 17.9 | **<0.001** | 9.1 | **<0.001** | 5.6 | 0.060<sup>†</sup> |
| Paternal population | 0.5 | 0.635 | 2.4 | 0.097 | 6.0 | **0.049** |
| Mat. Pop * Pat. Pop | 0.2 | 0.840 | 0.9 | 0.437 | 14.5 | **0.006** |
| Block | n/a | n/a | 4.6 | **0.001** | 65.3 | **<0.001** |
| Contrast | *t* | *P* | *t* | *P* | $\chi^2$ | *P* |
| BET<sub>pool</sub> VS. WIN<sub>pool</sub> | n/a | n/a | 0.3 | 0.736 | 5.7 | **0.017** |
| Fou: BET vs. WIN<sub>pool</sub> | n/a | n/a | 3.1 | **0.002** | 21.4 | **<0.001** |
| Tip: BET vs. WIN<sub>pool</sub> | n/a | n/a | -1.0 | 0.312 | 0.7 | 0.408 |
| Ver: BET vs. WIN<sub>pool</sub> | n/a | n/a | -0.8 | 0.410 | 0.2 | 0.672 |
| Fou: BET vs. WIN | n/a | n/a | 1.8 | 0.079 | 30.6 | **<0.001** |
| Tip: BET vs. WIN | n/a | n/a | -1.3 | 0.195 | 0.0 | - |
| Ver: BET vs. WIN | n/a | n/a | 0.6 | 0.565 | 0.0 | - |

After stratification, we sowed seeds (5 per cell) into a 72-cell flat insert (cell volume = 70 ml, Greenhouse Megastore, Danville, Illinois, USA), filled with a 3:1 mixture of MetroMix potting soil (Sun Gro Horticulture, Agawam, Massachusetts, USA) and autoclaved sand. A total of 30-50 seeds were sown for each population-stratification treatment combination. The seeds were then germinated in an incubator (Percival scientific, Perry, Iowa, USA) set to a constant 27°C and a 12-hour photoperiod. After approximately 4 weeks, cells were thinned to one seedling per cell, and the flat was moved to a heated greenhouse (minimum: 20°C) with supplemental lighting (12-hour photoperiod). Plants were grown there for approximately 6 weeks, watering as needed. Plants were then moved to a large refrigerator (4°C) with lights (10-hour photoperiod) for approximately one month to synchronize flowering. After the end of this vernalization period in mid-January to late February (depending on stratification treatment) 2020, plants were moved back to the greenhouse and transplanted into 17 cm diameter circular pots (volume = 2.4 L, Greenhouse Megastore, Danville, Illinois, USA) filled with the same soil mixture. Plants were watered as needed and fertilized monthly with ¼ strength Hoagland’s solution.

Once plants had begun to flower (early March 2020), we performed controlled hand pollinations within-populations and between-populations in all possible combinations and crossing directions (Figure S1A). In mid-March, because of COVID related shutdowns, the plants (19, 13, and 10 individuals from Fou, Tip, and Ver respectively) were moved to a private residence and grown on light racks with a 14-hour photoperiod for the duration of the hand pollinations. Most of the pollinations were performed from mid-March through the end of June 2020. Before each pollination we identified ovule donor flowers several days prior to anthesis and removed the upper part of the corolla tube, filaments, and anthers with scissors and forceps. Pollinations were completed when the stigma lobes of the ovule donor flowers appeared to be receptive (usually 3-4 days later) by removing stamens from a pollen donor flower with forceps and brushing the dehisced anthers against the stigma. Each pollination was tracked with a unique color combination of embroidery floss tied around the pedicel. Pollinations were done haphazardly based on flower availability, but we attempted to balance the cross types done for a given maternal population on a given day to minimize confounding treatment effects with age effects. Because we started with bulked seed collections and were interested in population level estimates of heterosis, and because of practical constraints of the number of receptive flowers per day on a given plant, we did not execute a balanced crossing design at the level of individual plants. However, most individual plants participated as both ovule and pollen donors, with population mean averages of 5-8 flowers as ovule donors and 4-7 flowers as pollen donors for between population crosses per plant. For within population crosses, each individual plant contributed an average (by population) of 3-4 flowers as ovule donors and 2-3 flowers as pollen donors.

The number of hand pollinations per maternal population ranged from 39-58 (mean = 46) for within-population (WIN) crosses and 85-102 (mean = 91) for between-population (BET) crosses, for a grand total of 411 crosses. The approximately two-fold greater number of hand pollinations for between-population crosses is because each maternal population was crossed to two different paternal populations, and we wanted sufficient power to test for interactions between maternal and paternal populations.

In addition to hand pollinations, we performed control emasculations where flowers were emasculated but no pollen was applied to the stigma lobes. Of the 69 control emasculations, less than 3% (2 emasculations) resulted in a fruit, giving us confidence that the vast majority of the pollinations performed were successful crosses between the intended pollen and ovule donors.

### Early fitness components

We collected fruits of all successful pollinations (n = 238; Table S2). This species required an extended period between pollination and fruit maturity (mean = 56.8 days, SD =10.4). There were no patterns in fruit set or the rate of fruit development with respect to cross type, but fruits of Tip developed on average about 5 days faster than fruits from the other two populations. For each of 17-39 WIN fruits and 45-71 BET fruits per maternal population (Table S2), we counted the number of seeds per fruit, excluding small, shriveled seeds presumed to be inviable.

We selected a subset of the viable seeds for each population and cross type, prioritizing fruits with enough viable seeds for germination assays and the goal of proportionally balanced sample sizes. This subset included approximately 300 seeds each of the WIN and the two BET combinations for populations Tip and Ver and approximately double those numbers for Fou (Table S2). These seeds were cold stratified as previously described for a period of 7 weeks starting in mid-November of 2020. Seeds were then sown into 72 cell flat inserts filled with the same soil mix (5 flats in total) with 7-12 (mean = 9.96, SD = 0.34) seeds per cell in a randomized block design. The number of cells per flat varied by maternal population with an average of 6 and 12 for WIN and BET crosses respectively from Tip and Ver, and approximately double the sample size for each cross type from Fou (Table S2). We recorded germination daily for a period of 24 days starting in mid-January 2021. Most seeds germinated within the first week, and only 5 new germinants were recorded during the second half of the scoring period. For estimates of heterosis, we used mean percent germination per cell recorded at the end of the scoring period.

Once seeds had germinated and seedlings had two true leaves, they were transplanted into individual cells of 72 cell flat inserts and moved to the greenhouse. The seedlings were transplanted into an incomplete randomized block design with a total of 30 flats (blocks). The number of cells per flat varied by maternal population, with an average of 5-6 for WIN and 10 for BET crosses from Tip and Ver, and an average of 11 WIN and 24 BET crosses from Fou. Any seedlings that died within the first week were replaced with another seedling of the same population-cross type. This mortality was likely due to transplant shock, and these individuals were omitted from subsequent analyses. Once roots started to poke through the bottom of the cells, individuals were transplanted first into 6 cm square (volume = 250 ml, Greenhouse Megastore, Danville, Illinois, USA) pots filled with the same soil mixture. Plants were watered as needed and fertilized twice with ¼ strength Hoagland’s solution. Cumulative juvenile survival was recorded up to approximately seven weeks after stratification, excluding individuals that died within one week of the initial transplant. Surviving plants that were established in the 6 cm pots were then randomly divided into two experiments: one in the greenhouse and one in the field simulating a new restoration.

### Greenhouse experiment

In late March 2021, seedlings were transplanted into 11 cm square (volume = 1L, Greenhouse Megastore, Danville, Illinois, USA) pots filled with the same potting mix in an incomplete randomized block design consisting of 48 flats each containing 7-8 pots. Each maternal population cross type combination was represented by 1-5 plants in at least 19 different flats (blocks). The total size of the greenhouse experiment was 354 plants with sample sizes for between population crosses being approximately double that of within population crosses, and sample sizes for Fou being approximately double that of the other populations (Table S2). Plants were grown for approximately 18 months, until mid-October of 2022, which included two flowering periods. During this time plants were watered as needed and fertilized once with Osmocote timed release 15-9-12 fertilizer at ½ the recommended dose after the first flowering period was over.

For each plant in the greenhouse experiment we scored adult fitness as the cumulative number of flowers produced over ∼18 months (two flowering seasons) including zeros for plants that died prior to reproducing or that did not reproduce. We additionally quantified two components of this composite measure of adult fitness separately: combined probabilities of survival and reproduction (did a plant reproduce or not over the entire course of the experiment), and cumulative fecundity (total flower number over the entire course of the experiment) of reproducing plants (Figure S1D). We used flower number as our measure of fecundity in the greenhouse because the plants would be unlikely to set fruit in the absence of hummingbird pollinators.

### Field experiment

We established a field experiment to quantify adult fitness components under more realistic restoration conditions. The site was located on the property of Arbor America LLC in West Point, Indiana (Tippecanoe County) USA. This location is within the extant range of *S. regia,* and the field experiment was planted on Coloma Sand (Mixed, mesic Lamellic Udipsamments), a soil type and series similar to the well-drained sites where *S. regia* naturally occurs (ISEE Network, 2024). Our experimental plot was embedded in a turfgrass area previously treated to establish a prairie strip (approximately 3m x 150m). This entailed an application of Roundup in May 2020 to remove the turf, followed by controlled burning approximately 1 month later, and another application of Roundup after approximately 2 more months. Herbicide applications and controlled burns are typical of establishing grassland restorations and help to promote native forb establishment (Török et al., 2024). In early March 2021, the plot was hand seeded with a short stature, high-diversity prairie mix from Spence Restoration Nursery (Muncie, IN, USA).

In early April 2021, seedlings were transplanted into a randomized complete block design consisting of six blocks containing 120 plants each. Blocks were established in small area of the prairie strip in a 2 x 3 grid, with an approximately 1m between, and around the edges, of blocks. In total there were 720 plants, with approximately double the sample size of the greenhouse experiment for each maternal population-cross type combination (Table S2). Each block was a rectangular grid of 3x40 plants with 30 cm spacing between the plants. Plants were hand watered for the first week only, and plants that died during this first week were considered to have succumbed to transplant shock and were replaced. In July of both 2021 and 2022, other vegetation within 10 cm of the base of experimental plants was clipped once to facilitate locating plants for the post-reproductive censuses. While the conditions of the experimental plot closely mimic the conditions during the initial phase of a restoration, we acknowledge that this species would likely be sown as seeds in an actual restoration. Given the extremely low rates of field establishment from seed for this species (Menges, 1995) and these populations in particular (< 2%, C. Oakley, unpublished data), and the effort to create enough seeds via experimental crosses, starting these plants in the field as seedlings was a necessary compromise.

In late October 2021, we recorded whether each plant was still alive and had reproduced, and for plants that had reproduced we counted the number of fruits, which persist on the plants for many months. It was logistically infeasible to repeatedly visit the field site frequently enough to count the number of flowers as was done in the greenhouse. Only about 11% of all plants had reproduced in the first year. We re-censused the same fitness components in October 2022. We estimated adult fitness as the total number of fruits (sum over both years) produced per individual transplanted, including zeros for plants that died before reproducing or surviving plants that did not reproduce. We additionally calculated the combined (across both years) probabilities of survival and reproduction for each plant and cumulative fecundity (total number of fruits across both years) for each reproducing plant (Figure S1D).

### Statistical analyses and estimates of heterosis

We analyzed each fitness component using ANOVA. We tested for the effects of maternal population (Fou, Tip, and Ver), paternal population, and their interaction. For all fitness components except seed number per fruit from the initial crosses (where there was no blocking possible), we included a term for block in the analysis to account for micro-spatial variation within the experiment. All effects were treated as fixed effects because the limited number of populations and blocks preclude treating them as a truly random sample. Inspection of residuals indicated no major issues with normality or heteroscedasticity for many fitness components, and for these we used normal error distributions. Juvenile survival and the combined probability of survival and reproduction in both the greenhouse and field experiments required a generalized linear model with a binomial error distribution. For each model with an interaction between maternal and paternal populations or main effects for both maternal and paternal population (with a threshold of P < 0.1), we used linear contrasts to test *a priori* hypotheses (with a threshold of P < 0.05). We first contrasted BET crosses (averaged over all populations and crossing directions; BET_pool_) to WIN crosses (averaged over all populations; WIN_pool_) to test if there was significant overall heterosis (or outbreeding depression).

We then performed two different sets of contrasts to test for heterosis (or outbreeding depression) separately by maternal population. First, we contrasted BET crosses (averaged over both paternal populations) to WIN crosses for each maternal population (Figure S1B). This is the simplest approach to investigating heterosis within maternal populations. However, this approach may lead to misleading conclusions if progeny from WIN crosses from different populations have very different mean fitness. To illustrate this issue, consider a hypothetical example where population A has very low within population fitness and population B has very high within population fitness. With purely additive effects (i.e. no dominance complementation or heterosis) the offspring of the crosses between these two populations should have an average fitness equal to the mean fitness of the two parents, a.k.a. the midparent value (Lynch and Walsh, 1998; Demuth and Wade, 2006; Oakley et al., 2015a). Comparing BET crosses to WIN crosses for population A could lead to the mistaken interpretation of heterosis, and a similar mistaken interpretation of outbreeding depression could result for the comparison in population B. To test for heterosis or outbreeding depression within maternal populations while accounting for additive differences in mean population fitness we performed contrasts of BET crosses for a given maternal population (averaged over both paternal populations) to WIN crosses averaged over all populations (because all populations were crossed in all combinations; WIN_pool_; Figure S1C). In discussing the results, we primarily focus on these contrasts but point out cases where results for the two methods differ notably. All statistical analyses were performed in JMP Pro v. 16.1.0 (JMP, 1989-2021).

For each fitness component and composite estimate of fitness we calculated estimates of heterosis both overall, and separately for each population. Overall heterosis (or outbreeding depression for negative values) was calculated as ([(BET_pool_-WIN_pool_)/WIN_pool_]*100), contrasting cross type means averaged over all populations. For estimates calculated separately by population, we used means for each maternal population-cross type combination averaging over the block means. For the simplistic approach we calculated heterosis as ([(BET-WIN)/WIN]*100). To account for differences in mean WIN fitness among populations to more accurately estimate non-additive effects, we averaged WIN crosses from all three populations ([(BET-WIN_pool_)/WIN_pool_]*100). We additionally calculated heterosis relative to the midparent value for each pair of maternal and paternal populations. However, relative fitness of BET crosses for different paternal populations were usually of the same sign (i.e. there was variation in the degree of heterosis or outbreeding depression for both populations, but never strongly contrasting directions of effects; Figure S2). We therefore focus on the estimates above where BET crosses were averaged across both paternal populations (Fig S1B and C).

For cumulative fitness, calculated separately for the greenhouse and field experiments, we used the product of four estimates: 1) seed number per fruit, 2) proportion germination, 3) proportion juvenile survival, and 4) adult fitness (Figure S1D). Adult fitness was calculated as total number of flowers per plant transplanted in the greenhouse, and total number of fruits per plant transplanted in the field, over the entire length of each experiment. Both measures of adult fitness include a combined probability of survival and reproduction and an estimate of fecundity, though the estimate of fecundity differs between the experiments (flower number in the greenhouse, fruit number in the field). Because this multiplicative measure of cumulative fitness yields only one estimate per cross type per population, we could not directly test the effects of population, cross type, and their interaction as with the fitness components. Other authors have approached this problem by using replicate line level estimates of multiplicative fitness (see for e.g., Dudash, 1990), but we were unable to use this approach both because of our initial bulked seed collection and because not all cross types were performed on all plants.

## RESULTS

### Early fitness components

For seed number per fruit and germination, there were significant effects of maternal population, indicating genetically based differentiation among maternal populations in mean values of these fitness components (Table 1). Averaging over all cross types, Fou had greater seed number per fruit and germination than the other two populations (Figure S3; Table S2). For seed number per fruit, there were no additional significant effects of paternal population or the interaction between maternal and paternal population, indicating no significant heterosis for this fitness component. For germination there was some indication of effects of paternal population, but there was no significant overall heterosis (BET_pool_ vs. WIN_pool_ contrast in Table 1). Pairwise contrasts of cross types separately by maternal population indicated significant heterosis (23%) for Fou (Figure 1; Tables 1&S4). For juvenile survival there was a significant interaction between maternal and paternal population (Table 1) and significant overall heterosis of about 6% (Tables 1&S3). Pairwise contrasts of cross types separately by maternal population indicated significant heterosis (13%) only for Fou (Figure 1, Tables 1&S4).

**Figure 1.**
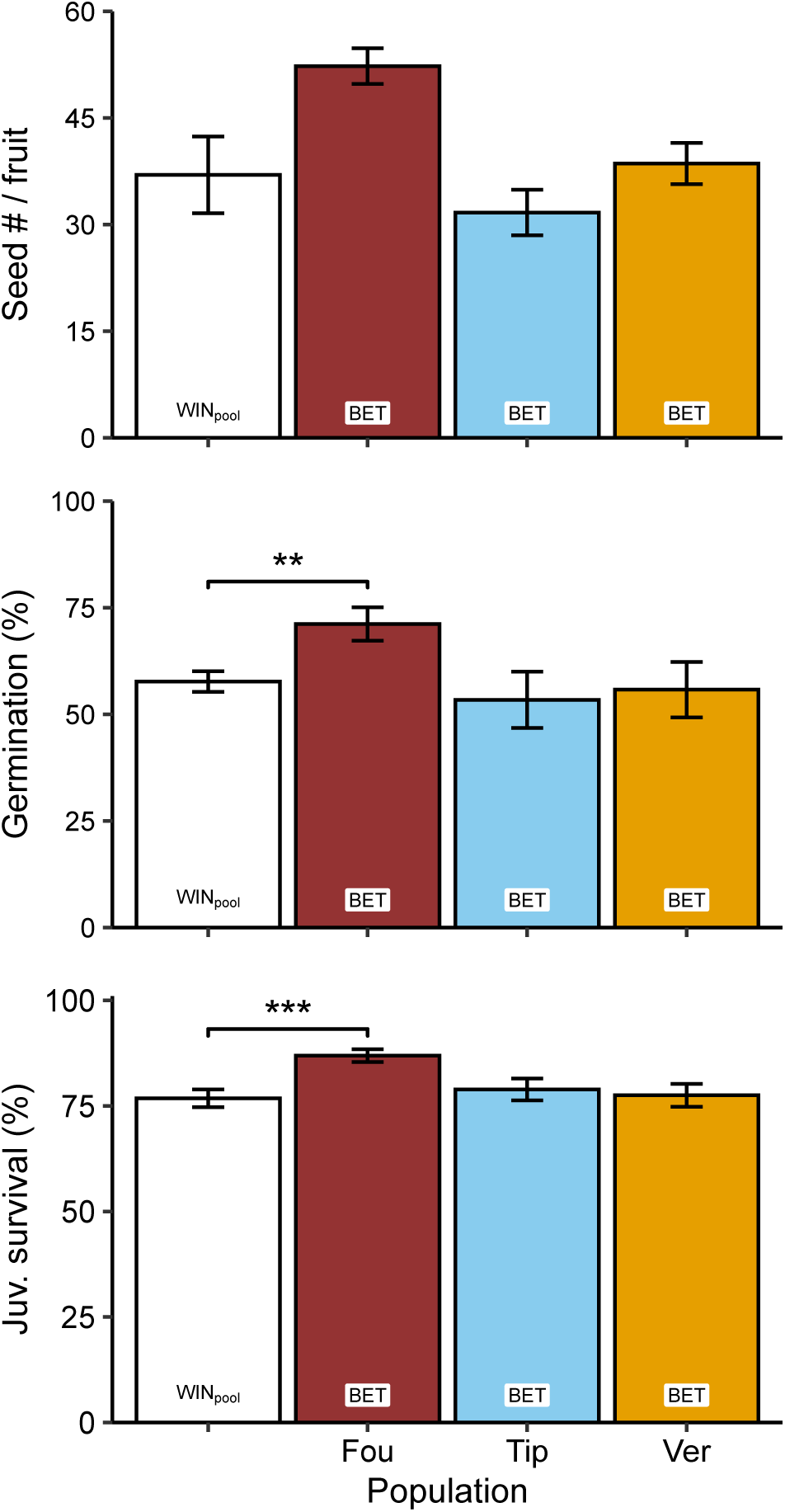
Mean fitness components following hand pollinations for each maternal population of *Silene regia* from western Indiana (Fou, Tip, and Ver) and cross type (WIN_pool_ = within-population crosses averaged over all three populations, BET = between-population crosses averaged over the two paternal populations) for seed number per fruit of the hand pollinations, germination, and juvenile survival. Means and standard errors were calculated from block means, except for seed number per fruit which are the raw means and SE. Sample sizes are given in Table S2. Significant contrasts of BET vs. WIN_pool_ are indicated where appropriate. ** P < 0.01, *** P < 0.001 (see Table 1).

### Greenhouse experiment

For adult fitness (flowers per seedling planted), there were no significant effects for any of the model terms (Figure 2; Table 2). This suggests a lack of genetically based differentiation for fitness, and the absence of heterosis in this environment. There were likewise no significant effects for the combined probability of survival and reproduction (Figure 2; Table 2). For fecundity, there was some indication of an interaction between maternal and paternal population (Table 2), with significant overall heterosis of about 19% (Tables 2&S3). Pairwise contrasts of cross types separately by maternal population indicated significant heterosis (36%) in Ver (Figs. 2&S4; Table S4). For this experiment, progeny from the three WIN crosses had similar mean fitnesses, and thus calculations of heterosis were qualitatively similar between the different approaches (Figs. 2&S4; Table S4).

**Figure 2.**
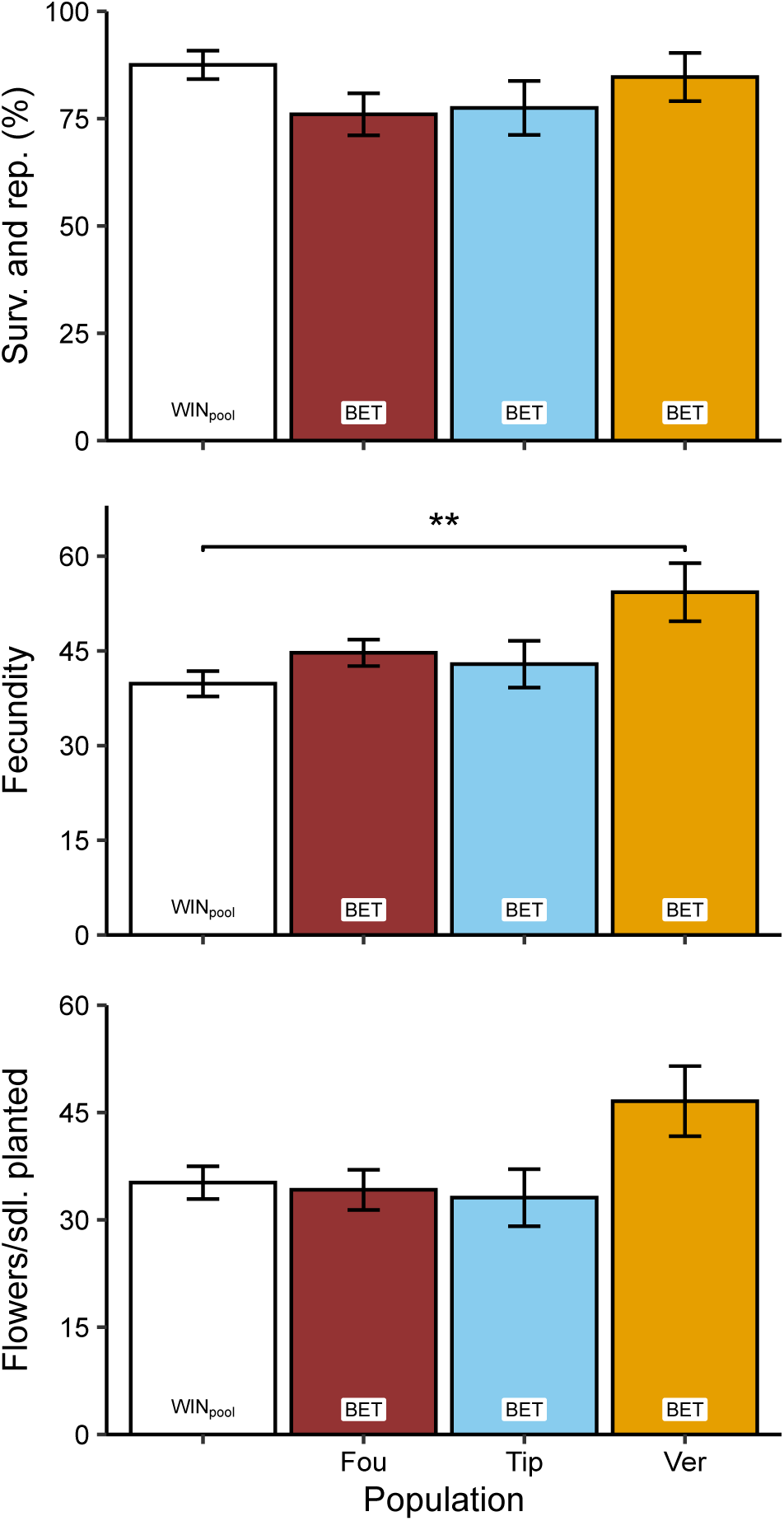
Mean fitness components following hand pollinations for each maternal population of *Silene regia* from western Indiana (Fou, Tip, and Ver) and cross type (WIN_pool_ = within-population crosses averaged over all three populations, BET = between-population crosses averaged over the two paternal populations) for the greenhouse experiment. Means and standard errors were calculated from the means within each of the 48 blocks. Significant contrasts of BET vs. WIN_pool_ are indicated where appropriate. ** P < 0.01 (see Table 2).

**Table 2.** ANOVA results in the greenhouse experiment for fitness components of offspring derived from hand pollinations within and between three populations (Fou, Tip, and Ver) of *Silene regia* from western Indiana. Fixed effects as in Table 1. The model for the combined probability of survival and reproduction employed a binomial error distribution, whereas the models for fecundity (flowers per reproducing plant) and flowers per seedling planted (including zeros) employed normal error distributions. Contrasts between cross types are only reported for models with a *P* value < 0.1 for either the interaction between maternal population and paternal population, or both main effects of maternal and paternal populations, otherwise these contrasts are n/a. For the contrasts, WIN = within-population cross, BET = between-population cross, and the subscript pool indicates that the contrast involves all individuals of a given cross type (averaged over all three populations), and contrasts that start with a population code followed by a colon test for heterosis or outbreeding depression separately by maternal population.

|  | Survival and reproduction |  | Fecundity (Denom. DF = 239) |  | Flowers per seedling planted (Denom. DF = 298) |  |
| --- | --- | --- | --- | --- | --- | --- |
| | $\chi^2$ | *P* | *F* | *P* | *F* | *P* |
| Maternal population | 0.8 | 0.675 | 1.4 | 0.250 | 1.6 | 0.194 |
| Paternal population | 1.9 | 0.380 | 0.8 | 0.459 | 1.6 | 0.208 |
| Mat. Pop * Pat. Pop | 3.5 | 0.483 | 2.1 | 0.080 | 1.1 | 0.372 |
| Block | 45.0 | 0.554 | 1.3 | 0.142 | 1.2 | 0.202 |
| Contrast | $\chi^2$ | *P* | *t* | *P* | *t* | *P* |
| BET<sub>pool</sub> vs. WIN<sub>pool</sub> | n/a | n/a | 2.5 | **0.015** | n/a | n/a |
| Fou: BET vs. WIN<sub>pool</sub> | n/a | n/a | 1.6 | 0.248 | n/a | n/a |
| Tip: BET vs. WIN<sub>pool</sub> | n/a | n/a | 1.2 | 0.235 | n/a | n/a |
| Ver: BET vs. WIN<sub>pool</sub> | n/a | n/a | 3.2 | **0.002** | n/a | n/a |
| Fou: BET vs. WIN | n/a | n/a | 1.0 | 0.300 | n/a | n/a |
| Tip: BET vs. WIN | n/a | n/a | 0.5 | 0.599 | n/a | n/a |
| Ver: BET vs. WIN | n/a | n/a | 2.5 | **0.014** | n/a | n/a |

### Field experiment

For adult fitness (fruits per seedling planted), there were significant effects of both maternal and paternal population (Table 3), indicating genetically based differentiation in mean fitness among populations. Most notably there was four-fold variation among maternal populations in mean fitness of progeny from WIN crosses, ranging from 2 fruits per seedling planted for Tip, to 3 for Fou, and 8 for Ver (Figure S5; Table S2). Additionally, there was a significant interaction between maternal and paternal populations which could indicate overall heterosis (or outbreeding depression) and/or variation among populations in the degree of heterosis (Table 3). Indeed, contrasts revealed significant overall heterosis (BET_pool_ vs. WIN_pool_) of about 45% (Tables 3&S3). For individual maternal populations, contrasts using the BET vs WIN_pool_ approach indicated significant heterosis (78%) only for Fou (Figure 3; Tables 3&S4), but at least on average, there was also heterosis for Tip (20%) and Ver (36%).

**Figure 3.**
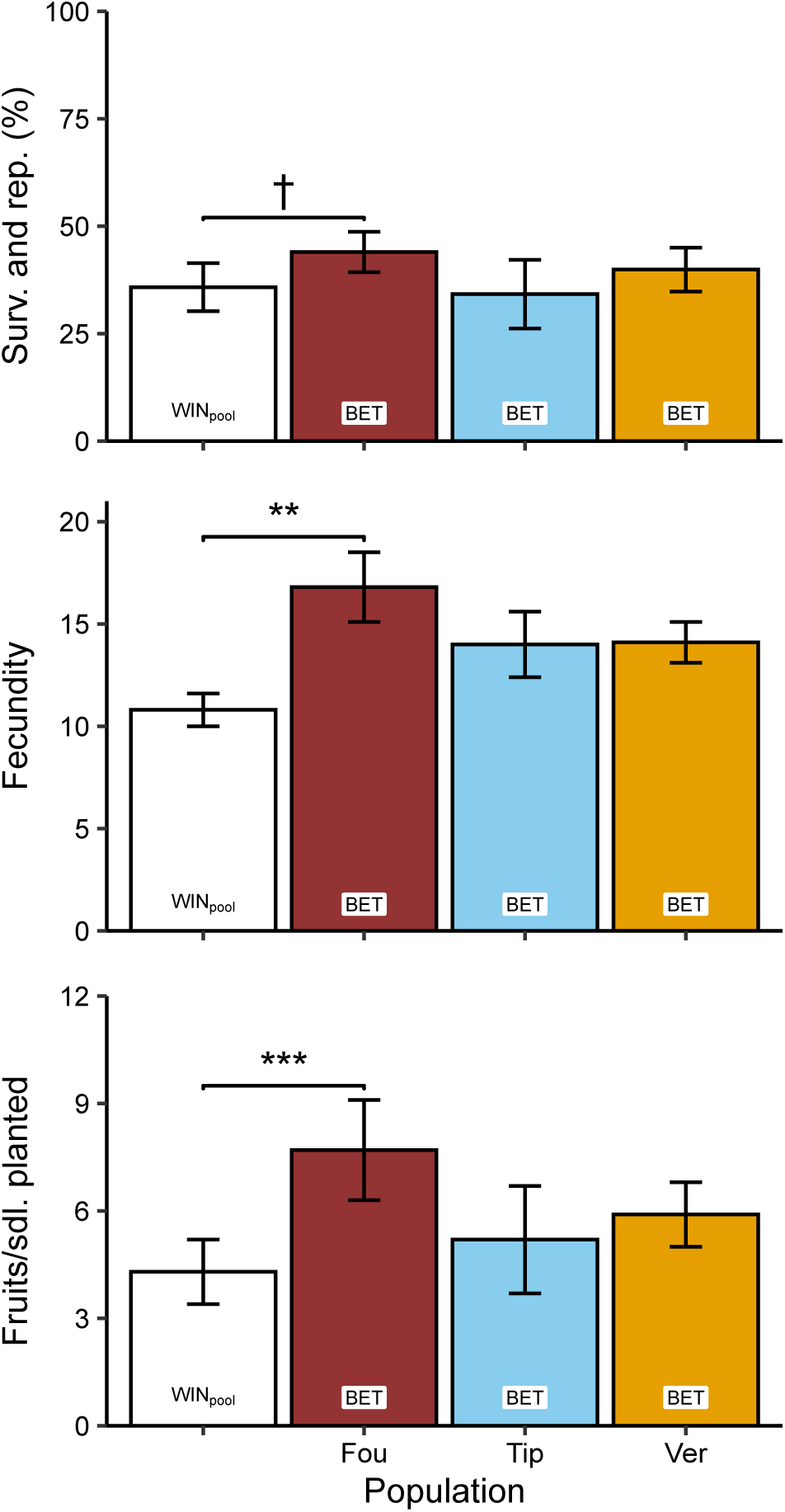
Mean fitness components following hand pollinations for each maternal population of *Silene regia* from western Indiana (Fou, Tip, and Ver) and cross type (WIN_pool_ = within-population crosses averaged over all three populations, BET = between-population crosses averaged over the two paternal populations) for the field experiment. Means and standard errors were calculated from the means within each of the 6 blocks. Significant contrasts of BET vs. WIN_pool_ are indicated where appropriate. † 0.10 > P > 0.05, ** P < 0.01, *** P < 0.001 (see Table 3).

**Table 3.**
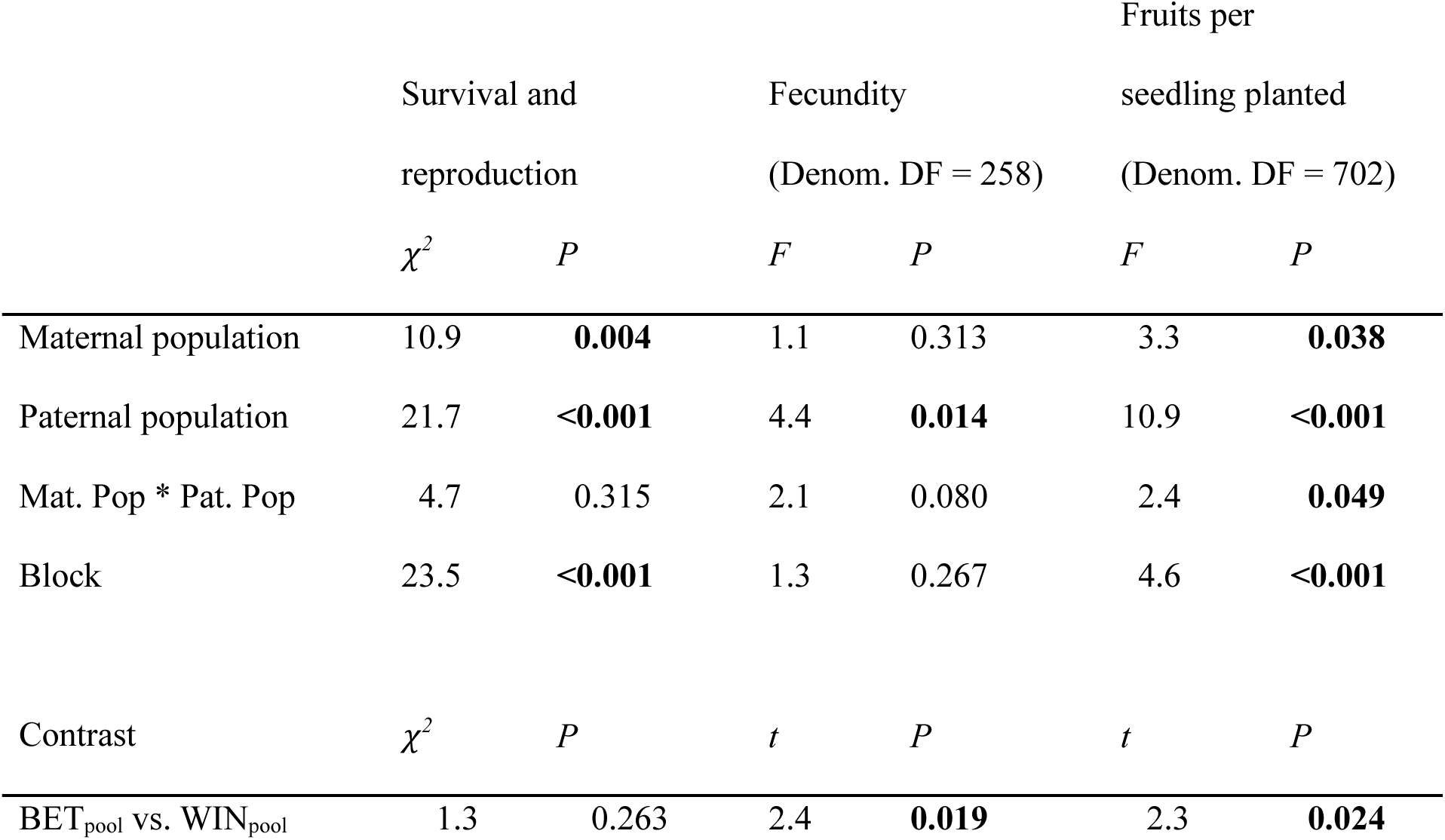
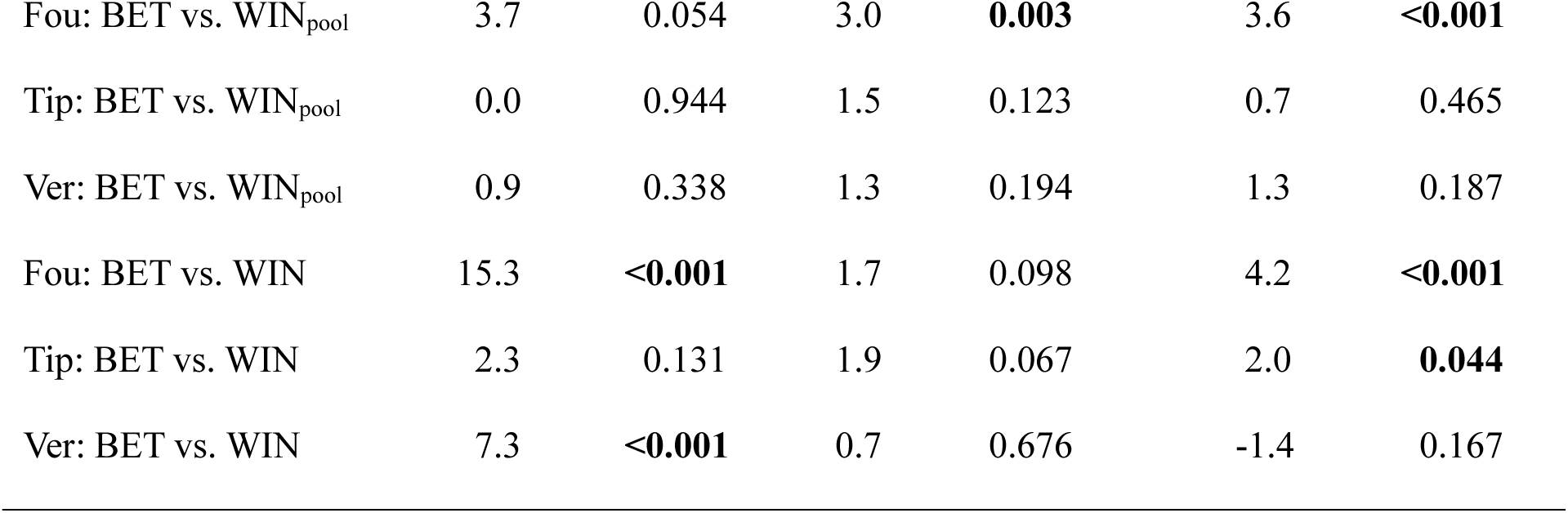
ANOVA results for the field experiment for fitness components of offspring derived from hand pollinations within and between three populations (Fou, Tip, and Ver) of *Silene regia* from western Indiana. Fixed effects as in Table 1. The model for the combined probability of survival and reproduction employed a binomial error distribution, whereas the models for fecundity (flowers per reproducing plant) and fruits per seedling planted (including zeros) employed normal error distributions. Contrasts between cross types are only reported for models with a *P* value < 0.1 for either the interaction between maternal population and paternal population, or both main effects of maternal and paternal populations, otherwise these contrasts are n/a. For the contrasts, WIN = within-population cross, BET = between-population cross, and the subscript pool indicates that the contrast involves all individuals of a given cross type (averaged over all three populations), and contrasts that start with a population code followed by a colon test for heterosis or outbreeding depression separately by maternal population.

|  | Survival and reproduction |  | Fecundity (Denom. DF = 258) |  | Fruits per seedling planted (Denom. DF = 702) |  |
| --- | --- | --- | --- | --- | --- | --- |
| | $\chi^2$ | *P* | *F* | *P* | *F* | *P* |
| Maternal population | 10.9 | **0.004** | 1.1 | 0.313 | 3.3 | **0.038** |
| Paternal population | 21.7 | **<0.001** | 4.4 | **0.014** | 10.9 | **<0.001** |
| Mat. Pop * Pat. Pop | 4.7 | 0.315 | 2.1 | 0.080 | 2.4 | **0.049** |
| Block | 23.5 | **<0.001** | 1.3 | 0.267 | 4.6 | **<0.001** |
| Contrast | $\chi^2$ | *P* | *t* | *P* | *t* | *P* |
| BET<sub>pool</sub> vs. WIN<sub>pool</sub> | 1.3 | 0.263 | 2.4 | **0.019** | 2.3 | **0.024** |
| Fou: BET vs. WIN<sub>pool</sub> | 3.7 | 0.054 | 3.0 | **0.003** | 3.6 | **<0.001** |
| Tip: BET vs. WIN<sub>pool</sub> | 0.0 | 0.944 | 1.5 | 0.123 | 0.7 | 0.465 |
| Ver: BET vs. WIN<sub>pool</sub> | 0.9 | 0.338 | 1.3 | 0.194 | 1.3 | 0.187 |
| Fou: BET vs. WIN | 15.3 | **<0.001** | 1.7 | 0.098 | 4.2 | **<0.001** |
| Tip: BET vs. WIN | 2.3 | 0.131 | 1.9 | 0.067 | 2.0 | **0.044** |
| Ver: BET vs. WIN | 7.3 | **<0.001** | 0.7 | 0.676 | -1.4 | 0.167 |

There was also significant variation among populations and population combinations for individual fitness components. For the combined probability of survival and reproduction there were significant effects of both maternal and paternal populations. As with adult fitness, the probability of surviving and reproducing for WIN crosses in Ver was much greater (about 1.6-fold) than for the other populations (Figure S5; Table S2). There was no significant overall heterosis (about 10% on average) for this fitness component (Tables 3&S3), and contrasts separately by maternal population suggest heterosis (23%) only in Fou (Figure 3; Tables 3&S4). For fecundity, there was a significant effect of paternal population and a suggestive effect for the interaction between maternal and paternal populations (Table 3). There was significant overall heterosis for fecundity of about 38% (Tables 3&S3). Contrasts separately by maternal population (Figure 3; Tables 3&S4) indicated significant heterosis (55%) only in Fou, but there was also heterosis for Tip (29%) and Ver (30%), at least on average.

Results for heterosis in the field experiment were notably different for some populations when fitness estimates of progeny from within population crosses were not averaged over all populations (Tables 3&S4). This is likely due to the extent of population differentiation in mean fitness estimates. For Tip, using the BET vs. WIN approach, we would conclude significant heterosis for adult fitness of 160% (compared to 20% above). The results for adult fitness for Ver do not qualitatively change (both non-significant) but there is a reversal in the sign of effects between the two approaches, with -26% outbreeding depression using the BET vs. WIN approach (compared to 36% heterosis above). For probability of survival and reproduction, the BET vs. WIN approach would indicate highly significant heterosis for Fou of 89% (compared to 23% above) and highly significant outbreeding depression of -34% for Ver (compared to 11% heterosis above).

### Cumulative fitness

Estimates of cumulative heterosis across all measured life stages were substantial in both greenhouse and field experiments, but they were generally stronger in the field experiment (Tables S3&S4). For the greenhouse experiment there was about 35% heterosis overall, and within populations heterosis was 92% for Fou and 35% for Ver, and there was weak (and non-significant based on results for fitness components) outbreeding depression (-23%) for Tip (Table S4). For the field experiment, overall heterosis was 112%. All three populations exhibited heterosis for cumulative fitness with considerable variation in magnitude among populations, from weak (6%) in Tip to modest (50%) in Ver to very strong (281%) in Fou (Table S4). Using the simplistic comparisons would cause the false conclusion of very strong heterosis (131%) in Tip and weak outbreeding depression (-4%) in Ver, but estimates for Fou were similar for both methods (Table S4).

## DISCUSSION

Seed sourcing strategies for restoration such as regional admixture provenancing have been suggested to help maintain both local adaptation and amounts of genetic variation within restored populations. Heterosis may provide an added short-term benefit to such strategies. To our knowledge, our study is the first to quantify heterosis for multiple fitness components in a newly established restoration. Averaged over all populations, the consequences of between population crosses for cumulative fitness was strongly positive (112% overall heterosis) in the field, suggesting that regional admixture provenancing should provide at least a short-term fitness advantage for this species. Estimates of heterosis from the field were greater than those from the greenhouse (35% overall heterosis). Additionally, life-stage specific estimates of both mean population fitness and heterosis varied among populations, particularly in the field experiment. This suggests that while there is an overall benefit of crosses between populations under realistic field conditions, relative fitness outcomes can depend on the populations involved and the environmental conditions of the fitness assay.

### Field restoration experiment: heterosis and variation in fitness among populations

Results from the field experiment simulating early restoration conditions are most directly relevant for restoration practitioners. Averaged over all of the populations there was significant heterosis for both fruits per seedling planted (45%) and fecundity (38%), and on average there were consistently beneficial effects on relative fitness for these measures (ranging from 20-78%) within populations. Heterosis for the combined probability of survival and reproduction was weaker (10% overall), and none of the results for individual populations were significant. When considering all life stages, there was no significant overall outbreeding depression for any fitness component. Thus, admixture of these populations in field restorations are likely to result in a net positive effect, at least for reproductive fitness, despite some among population variation in the degree of heterosis.

The among population variation in mean fitness in the field experiment is important for the restoration potential and the calculation and interpretation of heterosis for individual populations. For example, Ver had much greater survival and adult fitness of WIN crosses compared to other populations. The greater mean fitness within Ver may be because it is the largest of the three populations and was more recently reduced in size, therefore Ver may harbor fewer fixed deleterious recessive alleles. Indeed, the mean of progeny derived from crosses within Ver exceeded that of progeny from between population crosses over all populations for fruits per seedling planted (VerWIN = 8.0 vs BET_pool_ = 6.3). However, cumulative fitness of between population crosses was greater than mean cumulative fitness within Ver (VerWIN = 102 vs BET_pool_ = 139), and admixture may also increase genetic variation important for future adaptation beyond that directly related to heterosis. The low mean adult fitness and fitness components within Tip for the field experiment was also notable, especially because the origin of this population was geographically closest to the planting site. This was the smallest population and may therefore suffer the most from fixed deleterious alleles, representing a situation in which the most “local” population may not always be the “best” source material for a restoration (Pizza et al., 2023). Where possible, modern approaches based on genomic data can inform seed provenancing strategies (Breed et al., 2019; Rossetto et al., 2019; Aitken et al., 2024; Fahey et al., 2025). It is unclear the extent to which genomic data could predict differences in mean fitness among these populations. It may be possible in the near future to apply genomic approaches to this species because of the development of genomic resources in the genus (Karrenberg et al., 2026).

The variation we observed in mean fitness among populations highlights the importance of using within population crosses averaged across populations as the frame of reference in calculating heterosis to avoid confounding additive and non-additive effects (e.g., field results for Ver). Estimating heterosis as a proportional deviation from the mid-parental value is commonly used for crosses between two natural populations with predominantly selfing mating systems (Lynch and Walsh, 1998; Oakley et al., 2019; Clo et al., 2021), but it should be broadly applicable to other systems. Alternatively, studies where different population categories (e.g. small vs. large) are represented by many populations (Paland and Schmid, 2003; Oakley and Winn, 2012; Willi, 2013) or where each maternal population is crossed with several paternal populations (Weisenberger et al., 2014; Spigler et al., 2017; Thoen et al., 2025) serve a similar purpose. These approaches minimize the influence of individual populations with very high or very low mean within population fitness by averaging over many populations. The use of such approaches to minimize confounding additive and dominance effects can also be important in greenhouse studies if the parental populations differ in mean fitness (Rojas-Gutierrez et al., 2026).

Field experiments of heterosis are logistically challenging but are increasingly more common, and like our study are often based on transplanted seedlings (Fenster, 1991; Ouborg and Van Treuren, 1994; Wagenius et al., 2010; Weisenberger et al., 2014; Oakley et al., 2015b; Spigler et al., 2017; Oakley et al., 2019; Thoen et al., 2025). We limit our discussion to studies of heterosis in rare or recently fragmented species because of the expected context dependency of heterosis on population size, genetic distance between parents, and spatial scale of the field experiment. Our overall conclusions agree with an exceptional 14-year field study that found inter-population variation in heterosis and limited evidence for outbreeding depression of seven fragmented populations of *Echinacea angustifolia* (Thoen et al., 2025). Another study in the same species conducted near the Thoen et al. (2025) plot also found no evidence for outbreeding depression, but curiously did not find any heterosis (Wagenius et al., 2010). Thoen et al. (2025) attribute differences between the results of the two studies to environment dependent expression of heterosis. Generally speaking, other field studies in rare taxa also find heterosis and little evidence for outbreeding depression (Weisenberger et al., 2014), though the magnitude of heterosis seems to depend on population characteristics (Pickup et al., 2012) and temporal environmental variation (Maschinski et al., 2013).

### Comparing heterosis across environments and life stages

Comparisons between field- and greenhouse-based estimates of heterosis are rare in the literature, though some studies have indicated that heterosis may be stronger under more stressful conditions (Maschinski et al., 2013; Hahn and Rieseberg, 2017; Stojanova et al., 2021). Our results indicate that heterosis is greater in field compared to greenhouse conditions for fruits per seedling planted (adult fitness) and both of its component parts: combined probability of survival and reproduction, and fecundity. We also found stronger average heterosis in the field compared to greenhouse for cumulative fitness. Greater heterosis under presumably stronger and more variable selection pressures in the field compared to the greenhouse agrees with results from many, though not all, similar comparisons for inbreeding depression (Dudash, 1990; Fox and Reed, 2010; Cheptou and Donohue, 2011).

In addition to the comparison between field and greenhouse, we were interested in how heterosis might change across different life stages. If heterosis is caused by the complementation of many mildly deleterious alleles, we would expect to see greater heterosis in later life history stages or in estimates that integrate multiple fitness components (van Treuren et al., 1993; Willi et al., 2007; Escobar et al., 2008; Oakley et al., 2015b; Spigler et al., 2017; Perrier et al., 2020; Thoen et al., 2025) as is seen for inbreeding depression in highly selfing species (Husband and Schemske, 1996). In general, the results for heterosis over all populations follow such a pattern, with no significant heterosis for the earliest two fitness components, seed number per fruit and germination, and very modest heterosis for seedling survival. It is important to note that in our study all early components of fitness were measured in the greenhouse. Similar experiments are rarely started from seed (but see Luijten et al., 2002), likely due to low field germination rates (e.g. Thoen et al., 2025) as expected for *S. regia* (Menges, 1995). Strong heterosis for early fitness components has been reported for some species even in controlled environment conditions (Busch, 2006; Oakley et al., 2015b; Söderquist et al., 2020), but outbreeding depression for early fitness components is also possible (Sletvold et al., 2012). For later life stages in the greenhouse, there was stronger overall heterosis compared to earlier life stages only for fecundity (19%). Incorporating additional years of fecundity estimates would likely increase overall heterosis in both the greenhouse and field environments if patterns of heterosis for fecundity are repeated across years. Thus, our measures of adult fitness components may underestimate overall heterosis.

## CONCLUSIONS

The heterosis reported here for crosses between these small remnant populations in conditions representative of a newly established restoration strongly suggests that regional admixture provenancing would be a beneficial strategy for this species. The short-term fitness boost from heterosis is in addition to any potential long-term benefits of increased genetic variation necessary to adapt to environmental heterogeneity and climate change. We agree with previous authors that outbreeding depression is unlikely to have serious negative consequences for restoration and conservation efforts (Frankham, 2015; Ralls et al., 2018; Bell et al., 2019; Hoffmann et al., 2021; Jordan et al., 2023) if appropriate guidelines are followed (Frankham et al., 2011; Weeks et al., 2011). Even if there is some outbreeding depression in recombinant generations in this system, selection is expected to quickly remove deleterious combinations (Luijten et al., 2002; Erickson and Fenster, 2006; Aitkin and Whitlock, 2013; Hamilton and Miller, 2016). In future studies of heterosis, we suggest crossing designs and/or analytical approaches (i.e. comparisons to mid-parent value for paired population designs) to interpret dominance effects correctly accounting for differences in mean fitness between populations. Furthermore, we suggest that it is important to report mean values for each cross type for each population in addition to estimates of heterosis/outbreeding depression. Many open questions about heterosis remain, and we agree with Menges (2008) that restorations should be seen as experimental opportunities in addition to their other benefits.

## ACKNOWLEDGEMENTS

We thank P. Chidambaram, C. Gallick, G. Lee, M. Lopp, N. Ryan, T. Soto, and J. Will for assistance with the experiments. We also thank G. Pardillo and Arbor America for permission to conduct the field experiment on their property, and K. Jacobsen for agreeing to house the plants for crossing for nearly four months during the COVID shutdown. We thank M. Dudash, S. Mantel, S. Mills, A. Schnable and anonymous reviewers for helpful comments on versions of the manuscript. This work was funded in part by USDA Hatch grant 7000281 to CGO via Purdue College of Agriculture, a National Science Foundation grant DEB-2325338 to CGO, and a Biodiversity Research grant from the Indiana Native Plant Society to IAT.

## AUTHOR CONTRIBUTIONS

I.A.T. and C.G.O. conceived of and designed the study. B.E. provided the seeds and established the field plot. I.A.T., J.D.R.G., and C.G.O. conducted the experiment and collected the data. C.G.O., J.D.R.G. and I.A.T. analyzed the data. J.D.R.G. produced the figures. I.A.T. and C.G.O. drafted the manuscript. All authors contributed to revising the manuscript.

## DATA AVAILABILITY STATEMENT

Upon acceptance all raw data will be made available at the Purdue Public Data Archiving Repository (PURR).

## SUPPLEMENTAL MATERIAL

Additional Supporting Information may be found online in the Supporting Information section at the end of the article.

**Figure S1.**
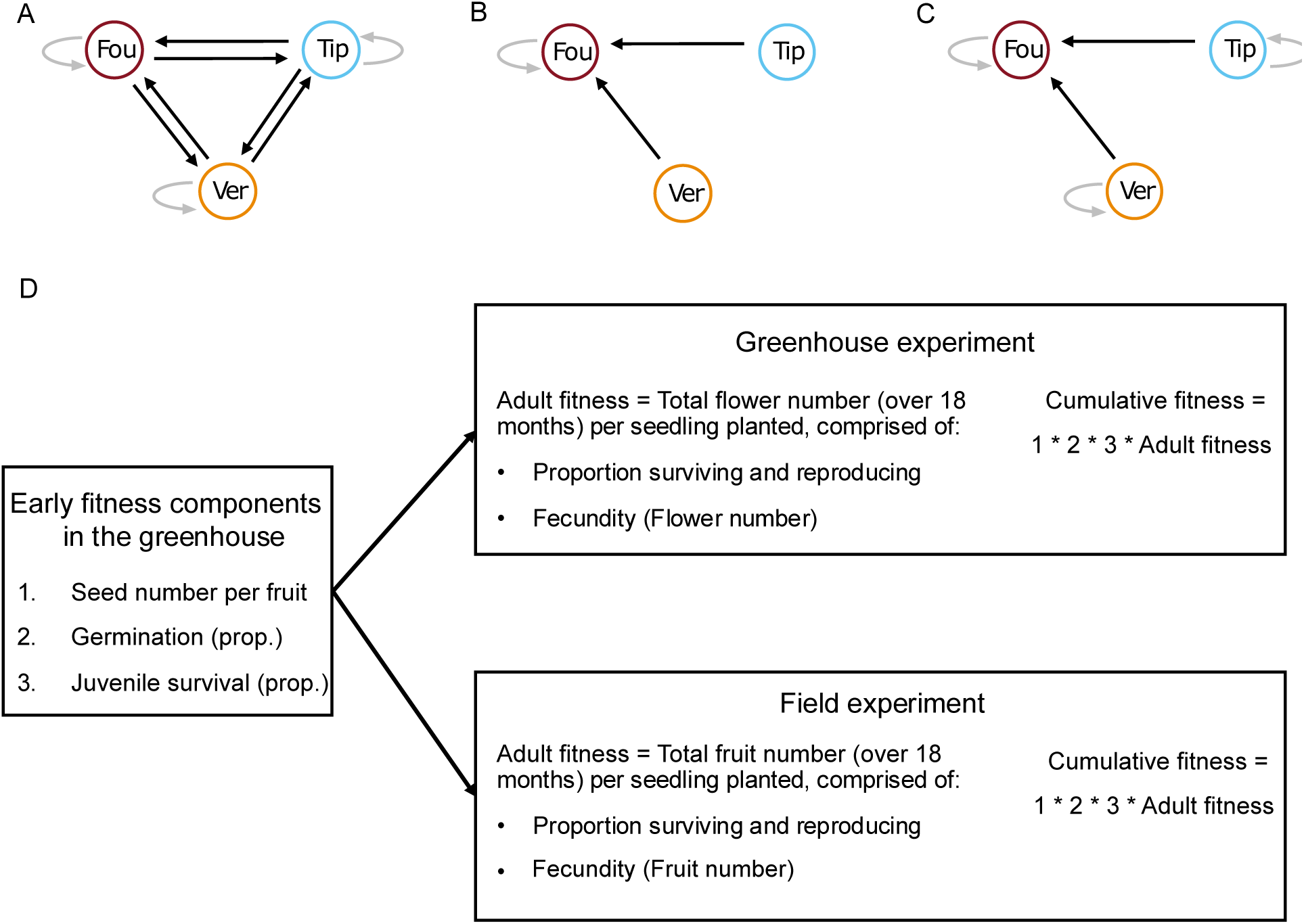
Schematic of crossing design, example comparisons of cross types, and list of fitness components and composite estimates of fitness in each experiment. The crossing design within (gray) and between (black) three populations of *Silene regia* from western Indiana (Fou, Tip, and Ver) are shown in A. For between population crosses the arrow points from the paternal population to the maternal population. B and C show the two different approaches for estimating heterosis using Fou as an example maternal population. B represents the simplistic approach where only the within population crosses for a given maternal population is used. In C, heterosis is measured as the increase in fitness of the mean of the between population crosses (averaged over the two paternal populations) relative to the mean of the within population (averaged over all three populations) crosses (WIN_pool_) to account for additive difference in mean fitness of within population crosses between the parents. D lists the fitness components measured in each experiment and indicates how cumulative fitness was estimated for the greenhouse and field experiments.

**Figure S2.**
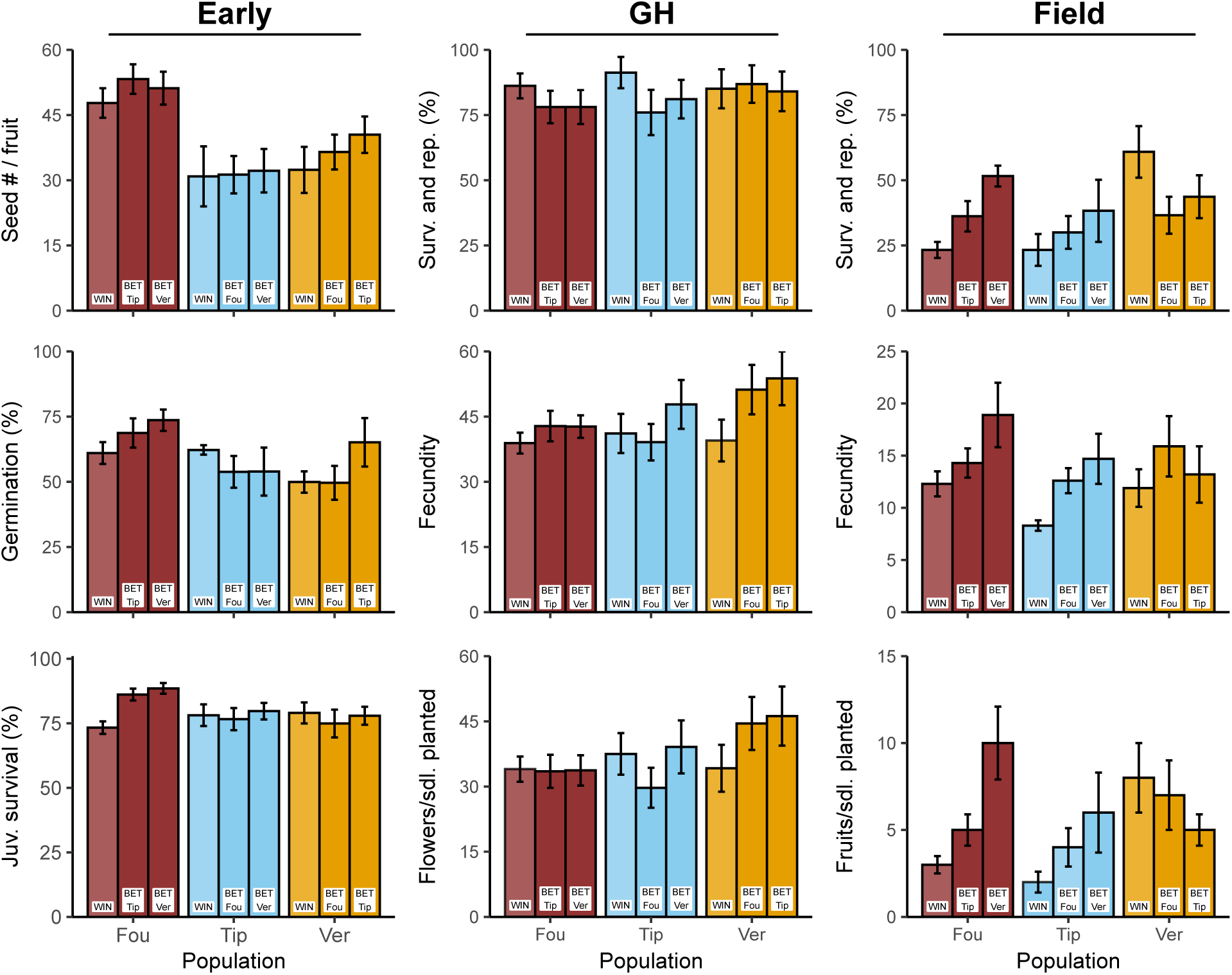
Mean fitness components following hand pollinations for each of the three maternal populations of *Silene regia* from western Indiana (Fou, Tip, and Ver) and cross type (WIN = within-population cross, BET = between-population cross) separately by the two paternal populations for between-population crosses. Means and standard errors were calculated from block means, except for seed number per fruit which are the raw means and SE. Sample sizes are given in Table S2.

**Figure S3.**
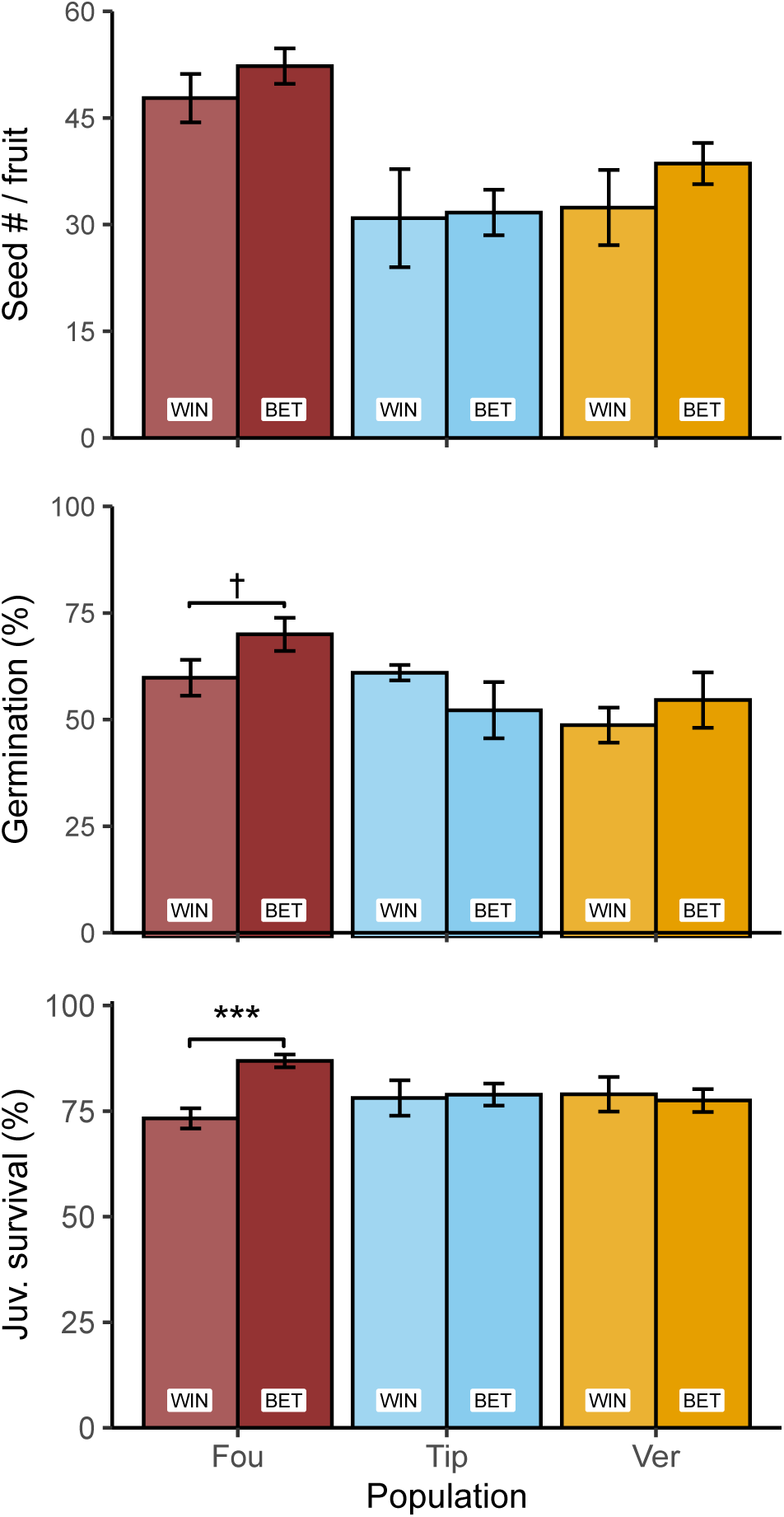
Mean fitness components following hand pollinations for each of the three maternal populations of *Silene regia* from western Indiana (Fou, Tip, and Ver) and cross type (WIN = within-population cross, BET = between-population cross) for seed number per fruit of the hand pollinations, germination, and juvenile survival. Means and standard errors were calculated from block means, except for seed number per fruit which are the raw means and SE. Sample sizes are given in Table S2. Within a population, significant contrasts of BET vs. WIN are indicated where appropriate. † 0.10 > P > 0.05, *** P < 0.001 (see Table 1).

**Figure S4.**
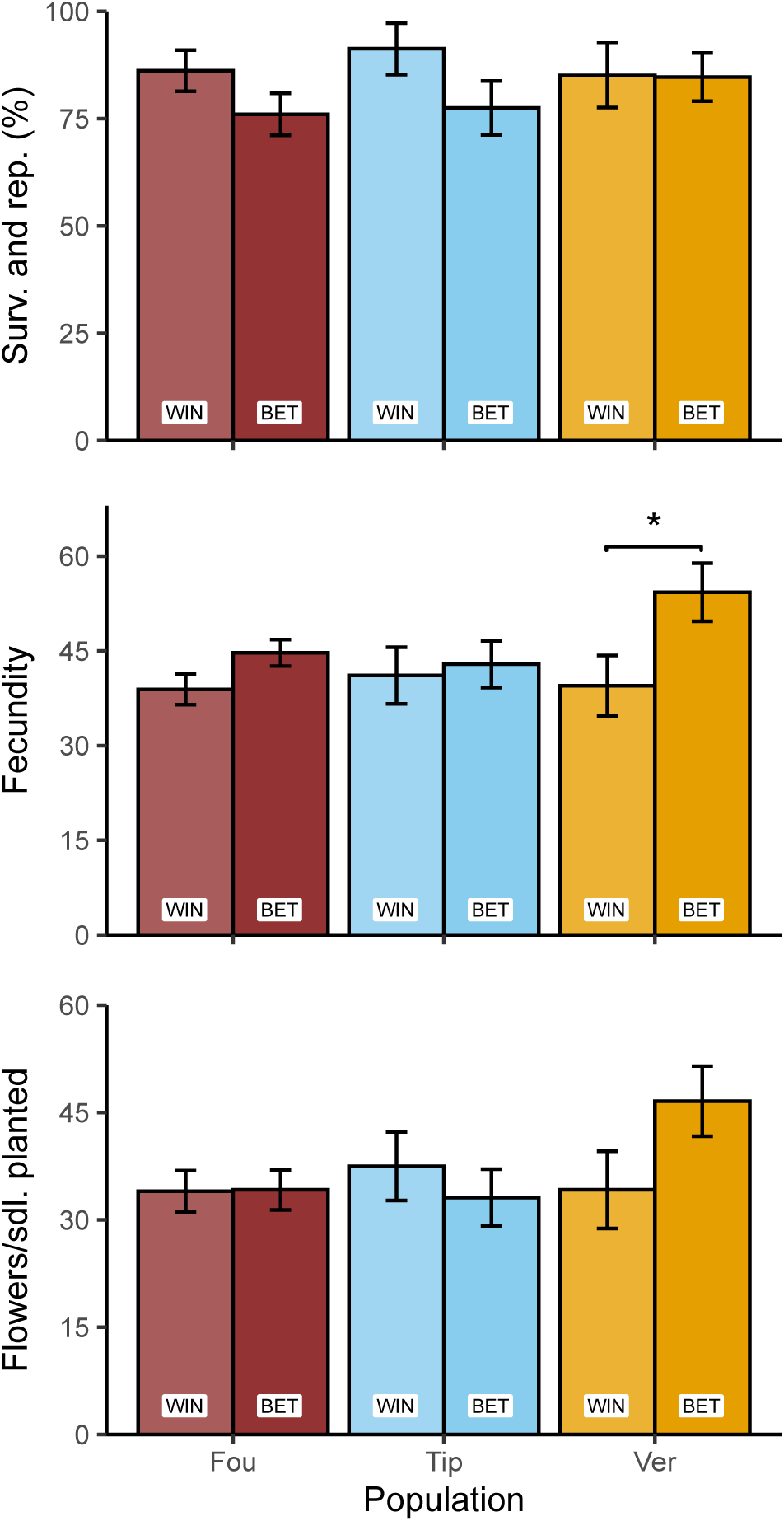
Mean fitness components following hand pollinations for each of the three maternal populations of *Silene regia* from western Indiana (Fou, Tip, and Ver) and cross type (WIN = within-population cross, BET = between-population cross) for the greenhouse experiment. Means and standard errors were calculated from the means within each of the 48 blocks. Within a population, significant contrasts of BET vs. WIN are indicated where appropriate. * *P* < 0.05 (see Table 2).

**Figure S5.**
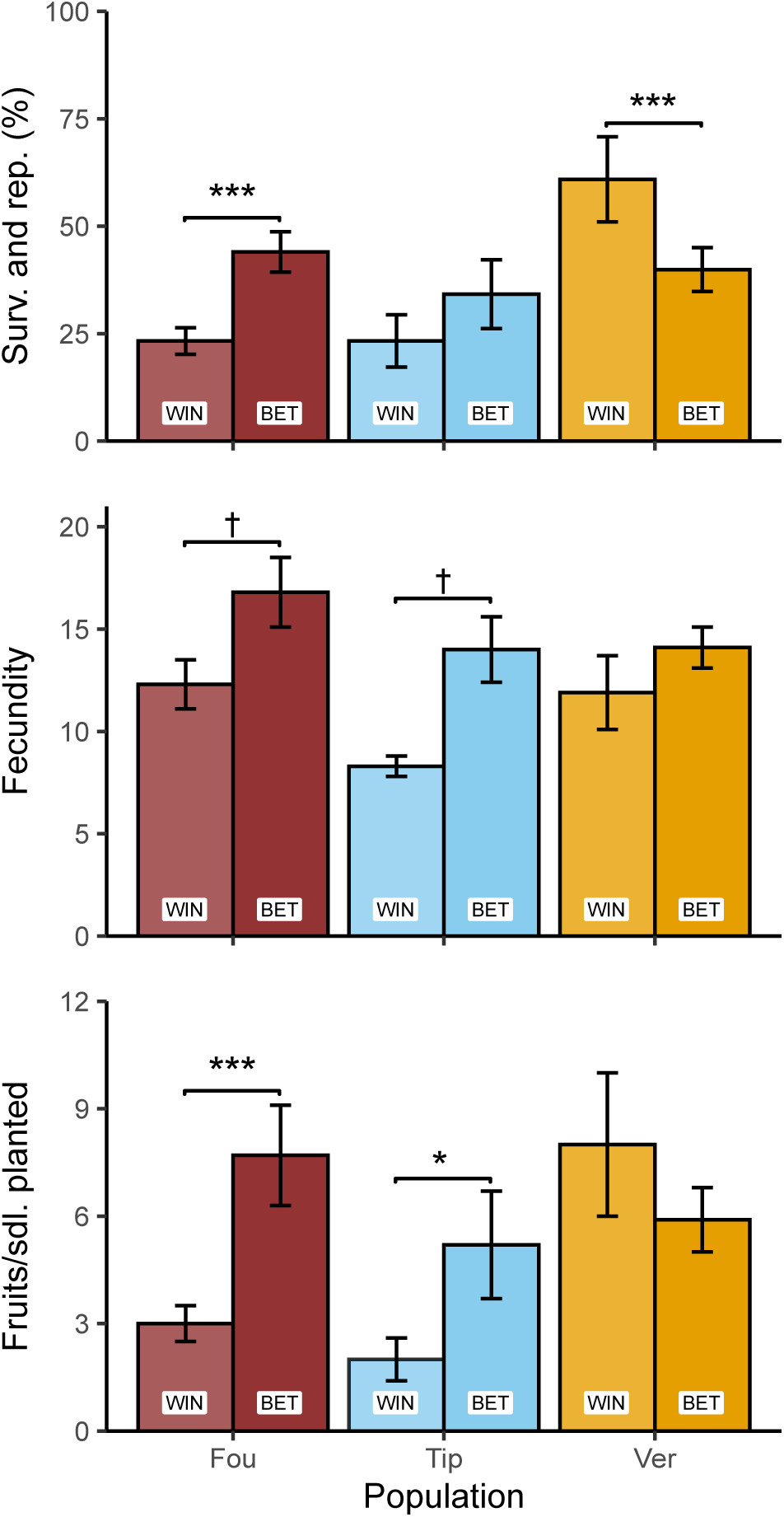
Mean fitness components following hand pollinations for each of the three maternal populations of *Silene regia* from western Indiana (Fou, Tip, and Ver) and cross type (WIN = within-population cross, BET = between-population cross) for the field experiment. Means and standard errors were calculated from the means within each of the 6 blocks. Within a population, significant contrasts of BET vs. WIN are indicated where appropriate. † 0.10 > P > 0.05, * *P* < 0.05, *** *P* < 0.001 (see Table 3).

**Table S1.** ANOVA results for early fitness components of offspring derived from hand pollinations within and between three populations (Fou, Tip, and Ver) of *Silene regia* from western Indiana. Fixed effects include the effects of maternal stratification treatment (10, 30, or 60 days), maternal and paternal populations and their interaction, and block (where appropriate, otherwise n/a). Models for both seed number per fruit and percent germination (per cell) employed normal error distributions (denominator degrees of freedom in parentheses). The model for juvenile survival employed a binomial error distribution. Contrasts between cross types are only reported for models with a *P* value < 0.1 for either the interaction between maternal population and paternal population, or both maternal and paternal populations, otherwise these contrasts are n/a. For the contrasts, WIN = within-population cross, BET = between-population cross, and the subscript pool indicates that the contrast involves all individuals of a given cross type (averaged over all three populations), and contrasts that start with a population code followed by a colon test for heterosis or outbreeding depression separately by maternal population. Model results were similar to models without incorporating effects of maternal stratification treatment (compare with Table 1) so we proceed with the simpler model throughout the results.

|  | Seed number per fruit<br>(Denom. DF = 227) |  | Germination<br>(Denom. DF = 344) |  | Juvenile Survival |  |
| --- | --- | --- | --- | --- | --- | --- |
| | *F* | *P* | *F* | *P* | $\chi^2$ | *P* |
| Maternal population | 13.1 | **<0.001** | 6.4 | **0.002** | 5.3 | 0.070 |
| Paternal population | 1.0 | 0.366 | 2.7 | 0.071 | 6.3 | **0.044** |
| Mat. Pop * Pat. Pop | 0.3 | 0.877 | 1.1 | 0.345 | 14.6 | **0.006** |
| Mat. Strat. Trt. | 13.5 | **<0.001** | 2.0 | 0.132 | 0.4 | 0.804 |
| Block | - | - | 4.5 | **0.002** | 64.9 | **<0.001** |
| Contrast | *t* | *P* | *t* | *P* | $\chi^2$ | *P* |
| BET<sub>pool</sub> vs. WIN<sub>pool</sub> | n/a | n/a | 0.3 | 0.758 | 6.0 | **0.015** |
| Fou: BET vs. WIN<sub>pool</sub> | n/a | n/a | 2.8 | **0.005** | 21.5 | **<0.001** |
| Tip: BET vs. WIN<sub>pool</sub> | n/a | n/a | -1.0 | 0.305 | 0.5 | 0.458 |
| Ver: BET vs. WIN<sub>pool</sub> | n/a | n/a | -0.6 | 0.520 | 0.3 | 0.577 |
| Fou: BET vs. WIN | n/a | n/a | 1.9 | 0.064 | 30.9 | **<0.001** |
| Tip: BET vs. WIN | n/a | n/a | -1.5 | 0.142 | 0.0 | - |
| Ver: BET vs. WIN | n/a | n/a | 0.6 | 0.525 | 0.0 | - |

**Table S2.** Sample sizes and mean values (averaged over flats/blocks where appropriate) for fitness components of offspring derived from hand pollinations within and between three populations (Fou, Tip, and Ver) of *Silene regia* from western Indiana. For seed number per fruit, sample size indicates the number of fruits. For germination, mean is the percent germination per cell (usually starting with 10 seeds) averaged over five flats, and sample size represents the total number of cells over all flats. For juvenile survival, the mean percent survival over the means of the 30 flats is given, and sample size represents the total number of seedlings over all flats. For the greenhouse experiment, sample sizes are the total number of plants at the start of the experiment over all 48 flats, and means are the mean over flat means. For the field experiment, sample sizes are the total number of plants at the start of the experiment over all 6 blocks, and means are the mean over block means. For fecundity in both experiments, sample sizes are the number of plants that reproduced. Adult fitness in the greenhouse is estimated by the total number of flowers per plant transplanted, and adult fitness in the field is estimated by the total number of fruits produced over the entire experiment per plant transplanted. Cumulative fitness was calculated separately for the greenhouse and field experiments as the product of four estimates: 1) seed number per fruit, 2) proportion germination, 3) proportion juvenile survival, and 4) adult fitness. No sample size (n/a) is given for cumulative fitness because it is the product of multiple fitness components.

|  | Fou |  |  |  | Tip |  |  |  | Ver |  |  |  |
| --- | --- | --- | --- | --- | --- | --- | --- | --- | --- | --- | --- | --- |
|  | BET |  | WIN |  | BET |  | WIN |  | BET |  | WIN |  |
|  | N | Mean | N | Mean | N | Mean | N | Mean | N | Mean | N | Mean |
| <u>Early</u> |  |  |  |  |  |  |  |  |  |  |  |  |
| Seed # / fruit | 71 | 52.3 | 39 | 47.8 | 45 | 31.7 | 17 | 30.9 | 47 | 38.6 | 19 | 32.4 |
| Germination (%) | 120 | 71.2 | 60 | 61.0 | 60 | 53.4 | 30 | 62.2 | 59 | 55.8 | 30 | 49.9 |
| Juv. survival (%) | 723 | 86.9 | 337 | 73.3 | 307 | 78.9 | 170 | 78.1 | 291 | 77.5 | 149 | 79.0 |
| <u>Greenhouse</u> |  |  |  |  |  |  |  |  |  |  |  |  |
| Surv. and rep. (%) | 118 | 76.0 | 59 | 86.2 | 59 | 77.5 | 30 | 91.3 | 59 | 84.7 | 29 | 85.1 |
| Fecundity | 112 | 44.7 | 55 | 38.9 | 53 | 42.9 | 28 | 41.1 | 55 | 54.3 | 26 | 39.5 |
| Flowers/sdl. planted | 118 | 34.2 | 59 | 34.0 | 59 | 33.1 | 30 | 37.5 | 59 | 46.6 | 29 | 34.2 |
| <u>Field</u> |  |  |  |  |  |  |  |  |  |  |  |  |
| Surv. and rep. (%) | 239 | 44.0 | 120 | 23.3 | 120 | 34.2 | 60 | 23.3 | 118 | 39.9 | 59 | 60.9 |
| Fecundity | 105 | 16.8 | 28 | 12.3 | 41 | 14.0 | 14 | 8.3 | 47 | 14.1 | 36 | 11.9 |
| Fruits/sdl. planted | 239 | 7.7 | 120 | 3.0 | 120 | 5.2 | 60 | 2.0 | 118 | 5.9 | 59 | 8.0 |
| <u>Cumulative</u> |  |  |  |  |  |  |  |  |  |  |  |  |
| Greenhouse | n/a | 1106.7 | n/a | 726.7 | n/a | 442.1 | n/a | 562.9 | n/a | 777.9 | n/a | 436.8 |
| Field | n/a | 249.2 | n/a | 64.1 | n/a | 69.5 | n/a | 30.0 | n/a | 98.5 | n/a | 102.2 |

**Table S3.**
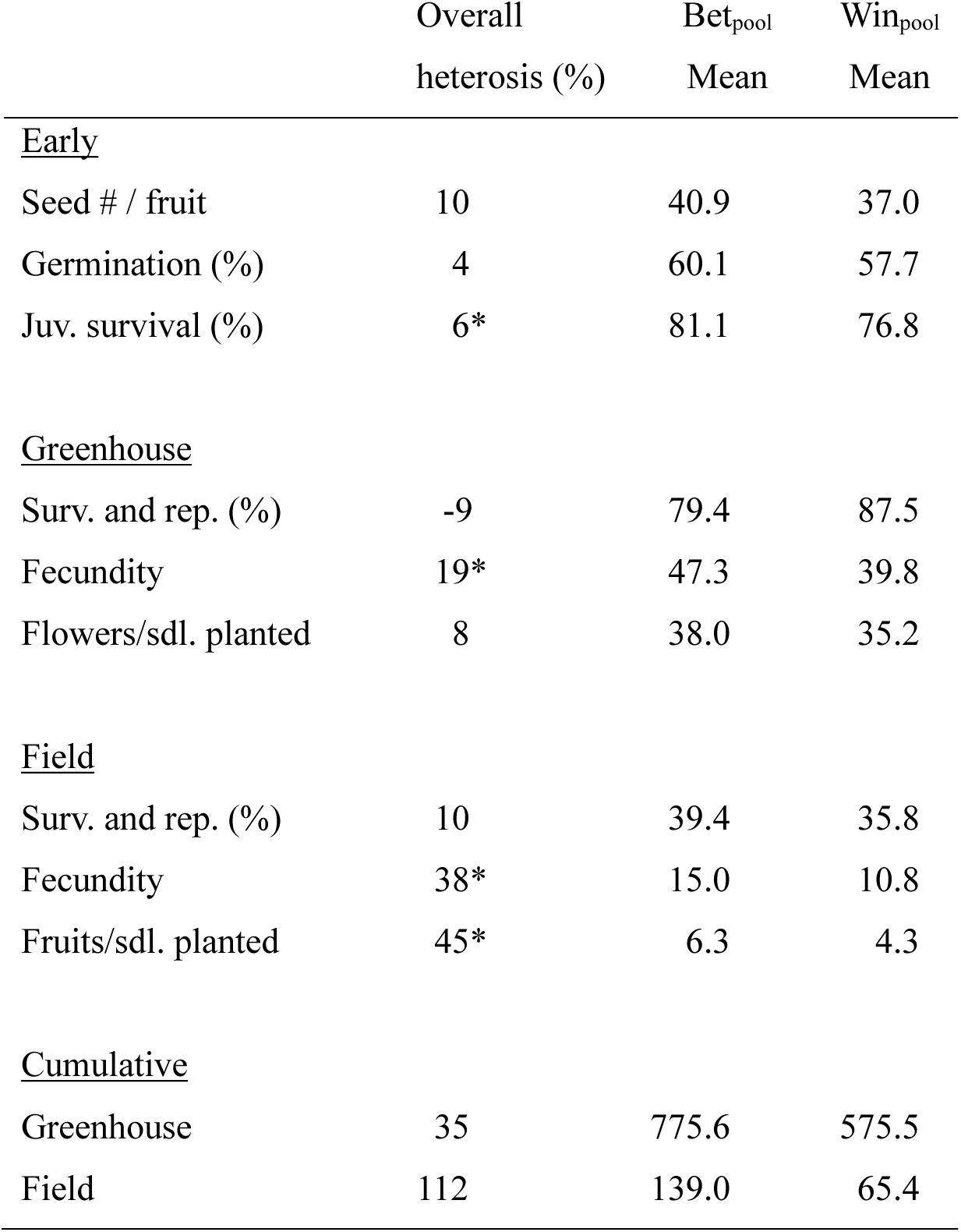
Estimates of overall heterosis (outbreeding depression for negative values) for fitness components and composite fitness estimates, averaging over all three maternal populations (Fou, Tip, and Ver) of *Silene regia* from western Indiana. Means for fitness components and composite estimates of fitness for each cross type, averaged over population cross type means (see Table S2), are also presented. Adult fitness in the greenhouse is estimated by the total number of flowers per plant transplanted, and adult fitness in the field is estimated by the total number of fruits produced over the entire experiment per plant transplanted. Cumulative fitness was calculated separately for the greenhouse and field experiments as the product of four estimates: 1) seed number per fruit, 2) proportion germination, 3) proportion juvenile survival, and 4) adult fitness. Significant contrasts corresponding to estimates of heterosis are indicated where appropriate. * *P* < 0.05. Significance testing of cumulative fitness was not possible (see methods).

**Table S4.**
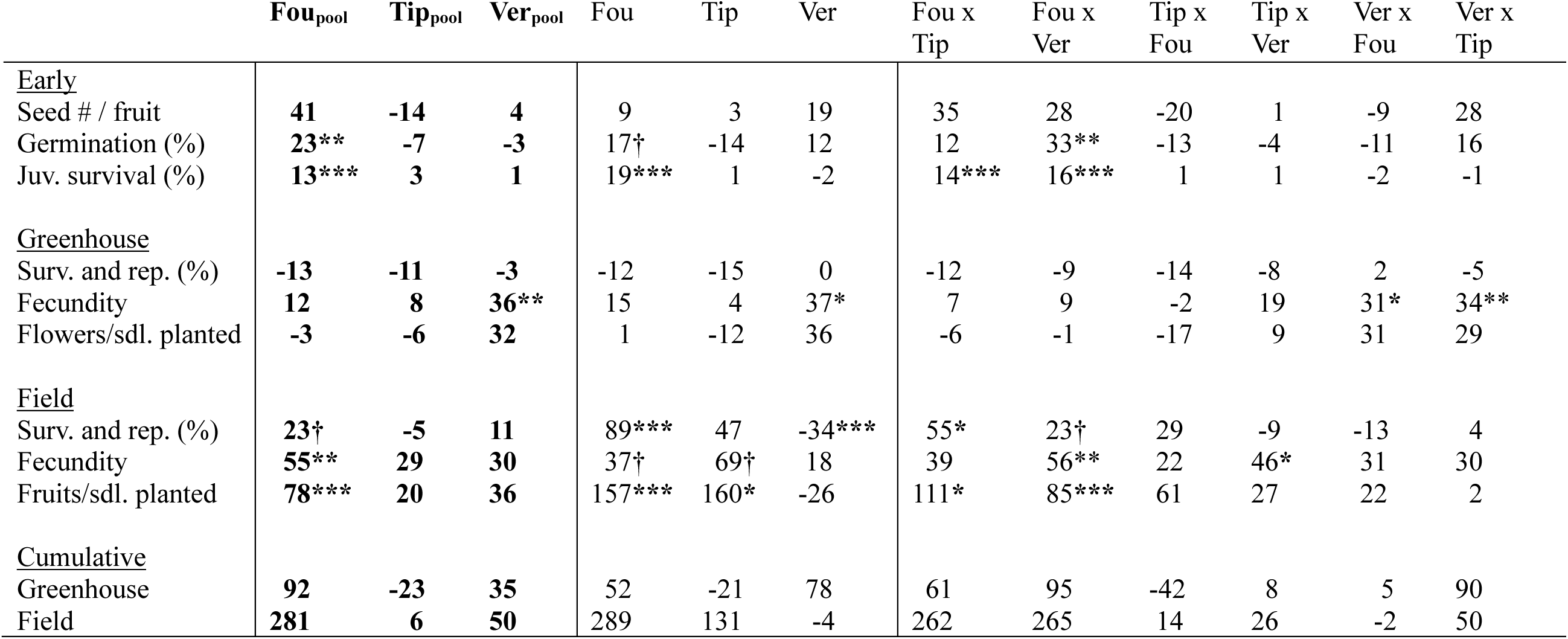
Estimates of heterosis (outbreeding depression for negative values) for each of the three maternal populations (Fou, Tip, and Ver) of *Silene regia* from western Indiana for fitness components and composite fitness estimates. Adult fitness in the greenhouse is estimated by the total number of flowers per plant transplanted, and adult fitness in the field is estimated by the total number of fruits produced over the entire experiment per plant transplanted. Cumulative fitness was calculated separately for the greenhouse and field experiments as the product of four estimates: 1) seed number per fruit, 2) proportion germination, 3) proportion juvenile survival, and 4) adult fitness. The first three columns are values for the averaging over all three populations for within-population cross approach used for the main results. The next three columns give values for the comparison of within- versus between- population crosses separately by maternal population. The remaining columns give the values for between-population values relative to the average of the within-population values for every possible combination of two populations in each direction (maternal x paternal). Within each set of columns significant contrasts are indicated as follows † 0.10 > P > 0.05, * *P* < 0.05, ** *P* < 0.01, *** *P* < 0.001. Significance testing of cumulative fitness was not possible (see methods).

|  | **Fou<sub>pool</sub>** | **Tip<sub>pool</sub>** | **Ver<sub>pool</sub>** | Fou | Tip | Ver | Fou x<br>Tip | Fou x<br>Ver | Tip x<br>Fou | Tip x<br>Ver | Ver x<br>Fou | Ver x<br>Tip |
| --- | --- | --- | --- | --- | --- | --- | --- | --- | --- | --- | --- | --- |
| <u>Early</u> |  |  |  |  |  |  |  |  |  |  |  |  |
| Seed # / fruit | **41** | **-14** | **4** | 9 | 3 | 19 | 35 | 28 | -20 | 1 | -9 | 28 |
| Germination (%) | **23\*\*** | **-7** | **-3** | 17† | -14 | 12 | 12 | 33** | -13 | -4 | -11 | 16 |
| Juv. survival (%) | **13\*\*\*** | **3** | **1** | 19*** | 1 | -2 | 14*** | 16*** | 1 | 1 | -2 | -1 |
| <u>Greenhouse</u> |  |  |  |  |  |  |  |  |  |  |  |  |
| Surv. and rep. (%) | **-13** | **-11** | **-3** | -12 | -15 | 0 | -12 | -9 | -14 | -8 | 2 | -5 |
| Fecundity | **12** | **8** | **36\*\*** | 15 | 4 | 37* | 7 | 9 | -2 | 19 | 31* | 34** |
| Flowers/sdl. planted | **-3** | **-6** | **32** | 1 | -12 | 36 | -6 | -1 | -17 | 9 | 31 | 29 |
| <u>Field</u> |  |  |  |  |  |  |  |  |  |  |  |  |
| Surv. and rep. (%) | **23†** | **-5** | **11** | 89*** | 47 | -34*** | 55* | 23† | 29 | -9 | -13 | 4 |
| Fecundity | **55\*\*** | **29** | **30** | 37† | 69† | 18 | 39 | 56** | 22 | 46* | 31 | 30 |
| Fruits/sdl. planted | **78\*\*\*** | **20** | **36** | 157*** | 160* | -26 | 111* | 85*** | 61 | 27 | 22 | 2 |
| <u>Cumulative</u> |  |  |  |  |  |  |  |  |  |  |  |  |
| Greenhouse | **92** | **-23** | **35** | 52 | -21 | 78 | 61 | 95 | -42 | 8 | 5 | 90 |
| Field | **281** | **6** | **50** | 289 | 131 | -4 | 262 | 265 | 14 | 26 | -2 | 50 |

